# RAD51 paralogs regulate double strand break repair pathway choice by limiting Ku complex retention

**DOI:** 10.1101/282855

**Authors:** Fernando Mejías-Navarro, Daniel Gómez-Cabello, Pablo Huertas

**Affiliations:** Departamento de Genética, Universidad de Sevilla, 41080, Sevilla, Spain; Centro Andaluz de Biología Molecular y Medicina Regenerativa (CABIMER), 41092 Sevilla, Spain; Genome Integrity Unit, Danish Cancer Society Research Center, DK-2100 Copenhagen, Denmark

**Keywords:** RAD51 paralogs, Homologous recombination, NHEJ, DNA repair pathway choice

## Abstract

RAD51 paralogs are a group of conserved proteins in eukaryotes that are involved in the repair of DNA breaks at several levels. On one hand, they help the strand invasion step catalyzed by RAD51. Also, they play late roles in Holliday Junction metabolism. Here we uncover a new role of the RAD51 paralogs at an earlier event in the repair of broken chromosomes. All five RAD51 paralogs affect the balance between double strand break repair pathways. Specifically, they favor homology-mediated repair over non-homologous end-joining. Such role is independent of RAD51 or the checkpoint activity of these proteins. Moreover, it defines a novel control point of double strand break repair independent and subsequent to DNA-end resection initiation.

Mechanistically, RAD51 paralogs limit the retention of Ku80 at the sites of DNA breaks. Thus, our data extend the role of this family of proteins to the earliest event of double strand break repair.

## Introduction

DNA double strand breaks (DSBs) are a common occurrence in most organism genomes, mainly due to the action of endogenous or exogenous chemical or physical insults (1–3). In order to minimize their impact, all cell types have a variety of molecular mechanisms available to repair those breaks (4–7). According to the use of a homologous sequence during the repair process, such pathways are commonly divided in two groups: non-homologous end-joining (NHEJ; homology-independent; (5)) or homologous recombination (HR; homology-mediated; (4)). NHEJ consists in the simple ligation with little or not processing of the ends (5). However, HR is a more complex molecular mechanism that requires extensive processing of the broken DNA, the recognition of the homologous template, etc. Indeed, HR can be subdivided in different subpathways (4), each of them using the homologous sequence in a distinct and particular way.

A central protein during recombination is the conserved RecA (prokaryotes)/Rad51 (eukaryotes) (4, 8). Rad51 catalyze the strand invasion step during associated to all subtypes of HR bar SSA, an intramolecular recombination between direct repeats (4). In all eukaryotes, in addition to the canonical Rad51, other proteins with partial sequence homology can be found, the so-called Rad51-paralogs (9). In mammalian, this family of proteins has 5 mitotic members: RAD51B, RAD51C, RAD51D, XRCC2, XRCC3 (9, 10). At the protein sequence level, all five paralogs and RAD51 share between 20 and 30% of identity. They can bind DNA, and all have weak ATP-ase activity. Biochemical studies have identified two distinct complexes in the cell: RAD51B–RAD51C–RAD51D–XRCC2 (BCDX2) and RAD51C–XRCC3 (CX3) (11). Several studies have firmly demonstrated the participation of RAD51 paralogs during HR in different eukaryotes (12–21) The classical role of RAD51 paralogs is to help RAD51 during the strand invasion step of recombination (13). In addition to this early function of the paralogs in recombination, they play additional roles at later steps, specifically during HJ resolution. Indeed, BCDX2 and CX3 complexes can bind *in vitro* to HJ (22, 23) and the CX3 complex was found associated with the resolution activity in protein extracts (24).

Moreover, some RAD51 paralogs are known to play a more general role during the activation of the DNA Damage Response (DDR) (25). The DDR is a signaling cascade that coordinates all cellular metabolism upon the appearance of DSBs on the DNA (2, 3). Briefly, when DNA damage is sensed the kinases ATM and/or ATR are activated, initiating a complex response of protein post-translational modifications, recruitment of proteins to the sites of DNA damage, changes on the DNA metabolism (transcription, replication and repair), cell cycle regulation, etc. In that context, RAD51C is included on this vast network. Indeed, RAD51C recruitment at damaged DNA is influenced by the DDR, but it is also required for efficient checkpoint signaling (25). Interestingly, RAD51C involvement on the DDR is related with the appearance of ssDNA, a key intermediate of all HR subpathways. In fact, the processing of the ends to create ssDNA during DSB repair, the so-called DNA end resection, is the best-known regulator of the choice between HR and NHEJ (6, 26). Furthermore, RAD51C mediates XRCC3 phosphorylation by ATR in a process that requires ssDNA formation (27). Also, XRCC3 is involved in the intra-S-phase and G2/M checkpoint (27). Such effect of RAD51 paralogs in checkpoint activation is more clearly observed at low doses of DNA damage (25, 27).

Here we take advantage of a reporter specifically designed to characterize how the decision between DSB repair pathways is made (28). Our data unveil a new role of RAD51 paralogs at even earlier steps of DSB repair that is independent of RAD51. Depletion of any of the five RAD51 paralogs leads to an unbalance of the NHEJ/HR ratio towards an increase of the former. Such function is not related with checkpoint activation defects. Strikingly, the paralogs influence the choice between DSB repair pathways without strongly affecting DNA end resection, uncovering a novel control point for such election. Finally, we demonstrate that RAD51 paralogs hamper Ku80 retention to the sites of DSBs. Thus, RAD51B, RAD51C, RAD51D, XRCC2 and XRCC3 provide a molecular link along all DSB repair timeframe, from the early selection of repair choice, through DNA strand invasion until HJ resolution.

## Results

### RAD51 paralogs are involved in DSB repair pathway choice independently to RAD51

RAD51 paralogs have been implicated at early and late events of HR (9–22, 24, 29). To investigate if such proteins might be involved in DSB repair at an earlier step, namely the choice between HR and NHEJ, we took advantage of the SeeSaw Reporter (28)(Figure 1A). Briefly, upon creating a DSB using the nuclease I-SceI, the reporter can be repaired by either NHEJ, maintaining an active GFP gene, or by a subtype of homologous recombination known as Single Strand Annealing (SSA), rendering an active RFP gene. So, the balance between NHEJ and HR can be calculated as the ratio between green versus red cells. The advantage of such system is that SSA does not require either strand invasion or HJ resolution. Thus, any phenotypes observed upon depletion of RAD51 paralogs cannot be accounted for the known roles in such recombination steps. Thus, we depleted RAD51B, RAD51C, RAD51D, XRCC2 and XRCC3 using shRNAs targeted against them (Figure S1). As controls, we used shRNAs against CtIP, RAD51 and a random sequence absent on the human genome (Scramble; Scr). As shown in Figure 1B, depletion of any of the five paralogs causes an unbalance of the NHEJ/HR towards an increase of NHEJ, in a similar fashion to the known regulator of DSB repair pathway choice CtIP. Although the result was not statistically significant upon RAD51C depletion, we think this was due to the high lethality observed in those cells. Lethality associated to the absence of all RAD51 paralogs is well documented in different model organisms (13, 30–35). In any case, as members of both the BCDX2 and CX3 complex showed similar results, we propose that both complexes must be involved. Moreover, and in agreement with the lack of strand invasion during SSA recombination, we concluded that such effect was independent on their role helping RAD51 to ssDNA, as it was not observed upon depletion of RAD51 itself. Thus, we postulate that RAD51 paralogs have a novel, RAD51-independent role on DSB repair at the level of choosing between HR and NHEJ. Specifically, we hypothesize that RAD51 paralogs are required to allow HR to outcompete with NHEJ.

**Figure 1.**
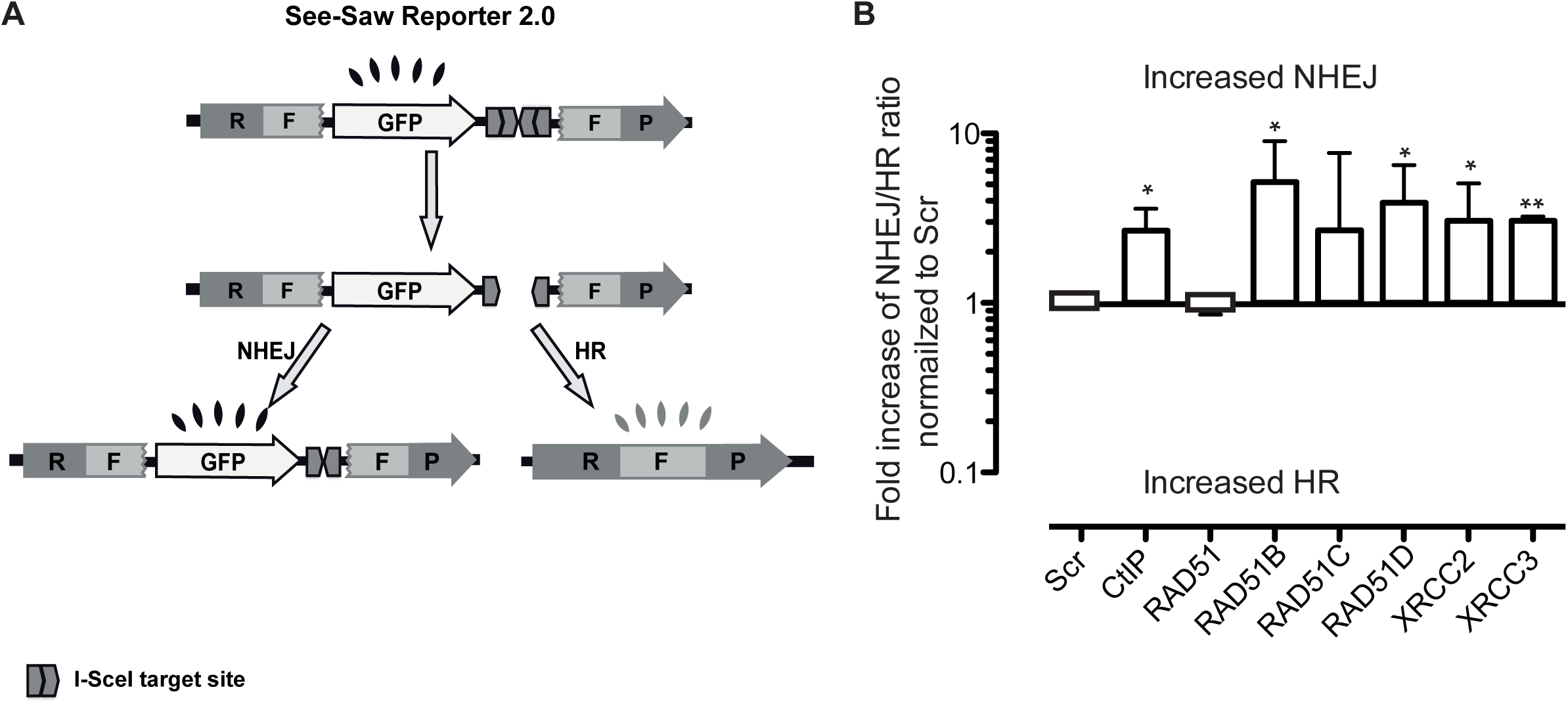
RAD51 paralogs affect DSB repair pathway choice. A, Schematic representation of the SeeSaw 2.0 reporter. A GFP gene is flanked by two truncated parts of RFP gene (RF and FP) sharing 302 bp of homologous sequence. Two I-SceI-target sites were cloned at the 3′end of the GFP gene in opposite orientation. After generation of a DSB by I-SceI expression, the damage may be resolved by NHEJ, thus cells will express the GFP protein, or using the homologous sequence by HR, creating a functional RFP gene. B, Effect of the depletion of CtIP, RAD51 and the RAD51 paralogs on NHEJ/HR ratio. To measure the deviation from the balance between NHEJ versus HR, the ratio between green versus red cells in each condition was calculated. To facilitate the comparison between experiments, this ratio was normalized with control cells transduced with an shRNA against a non-target sequence (Scr). Those conditions that skew the balance towards an increase NHEJ result in fold increase above 1. Data represent a minimum of three sets of duplicated experiments.

### RAD51 paralogs role in DSB repair pathway choice is independent of DNA-end resection

DSB repair pathway choice is tightly regulated. So far, there are two well-known, related, regulatory events: cell cycle and DNA-end resection. HR can only take place during the S and G2 phases of the cell cycle, whereas NHEJ takes place also in G1 (6, 26). This is due to the restriction of DNA-end resection to S and G2. Mechanistically, this is controlled by the activation of CtIP by CDK phosphorylation in S and G2 (36, 37), and by a crosstalk between two pair of protein duplexes: RIF1-53BP1 complex, which protects DNA ends from resection mainly in G1, and the BRCA1-CtIP complex, which promotes resection mainly in S and G2 (38, 39). In order to investigate how RAD51 paralogs control the choice between HR and NHEJ, we first analyzed if their depletion affect cell cycle distribution (Figure 2A). As it was not significantly impaired, we wondered if the number of cells that were resecting their DNA was influenced by RAD51 paralogs. Due to the high lethality observed upon depletion of some RAD51 paralogs, mainly RAD51C, we decided to focus in specific paralogs. We choose XRCC3, from the CX3 complex. From the BCDX2, as it has been proposed that there are two subcomplexes (BC and DX2) (11, 40), we selected RAD51B and XRCC2 to continue our experiments. Thus, we analyzed the recruitment upon DNA-damage of the anti-resection protein RIF1 and the ssDNA protecting complex RPA (Figures 2B-C) and found no differences that could explain the unbalance between HR and NHEJ upon depletion of RAD51 paralogs. One possibility was that, although cells were still initiating resection proficiently after infection with shRNAs targeted against RAD51 paralogs, resection rate was mildly impaired, i.e. the length of the resected DNA was shorter. This type of moderate resection phenotype has been observed previously upon depletion of BRCA1 (41). To test this idea, we took advantage of a high-resolution technique (Single Molecule Analysis of Resection Tracks; SMART) that measures the length of resected DNA in different experimental situations (41, 42). With this high-resolution technique, we could observe only a small effect on DNA end resection when compared with control cells (Figure 2D). Indeed, only XRCC2 depleted cells showed a significant shortening of resected DNA.

**Figure 2.**
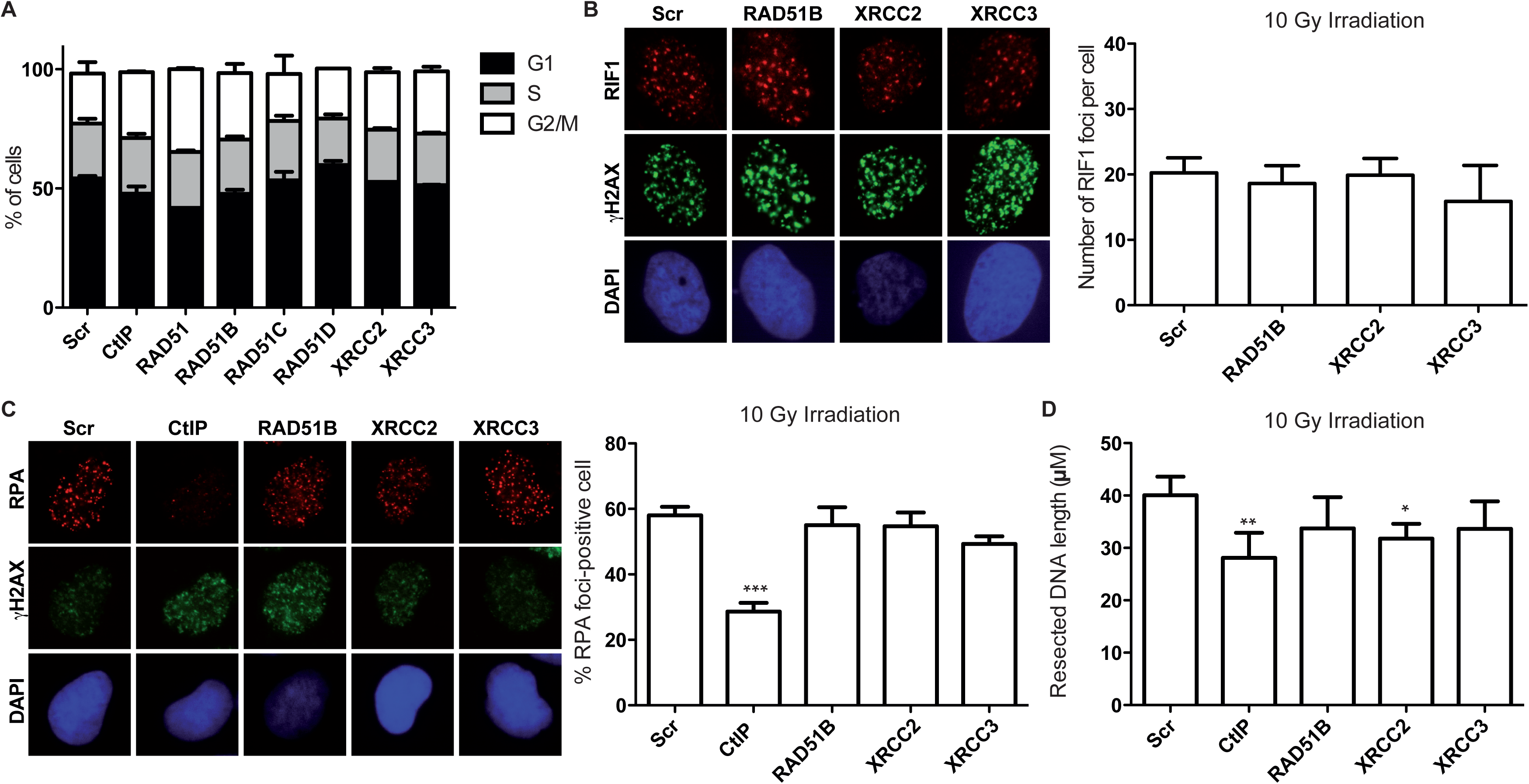
RAD51 paralogs do not affect DNA end processing. A, Cell cycle distribution of U2OS cells transduced with shRNAs targeted against the indicated genes. For details see methods section. B, Cells depleted of the indicated proteins were irradiated (10 Gy) and let to recover for 1 h. A representative image of RIF1 foci upon each gene downregulation is shown. The number of foci per cell was calculated in at least 100 cells. Graph represents the average of the median and standard deviation of three independent experiments. C, Cells were treated as described in (B) and immunostained with an anti RPA32 antibody. A representative image of RPA foci upon depletion of each protein is shown. The number of cells with visible RPA foci was scored. Other details as in B. D, Cells were irradiated and let to recover for 1 h before the length of resected DNA was calculated as described in the methods section. The average and standard deviation of three independent experiments is shown.

### RAD51 paralogs regulation of the NHEJ/HR ratio is not related with checkpoint activation

Checkpoint activation regulates the global response to DNA damage. Using the SSR, we have previously shown that checkpoint is required to control DSB repair pathway choice (28). As several RAD51 paralogs are involved in checkpoint activation at low damage doses (25, 27), we investigate if that was how they regulate the NHEJ/HR ratio. We tested checkpoint activation upon depletion of RAD51B, XRCC2 and XRCC3 at different doses of DNA damage. As shown in figure 3, we could not observe any effect on checkpoint activity in our experimental setup, neither at high nor at low doses of irradiation.

**Figure 3.**
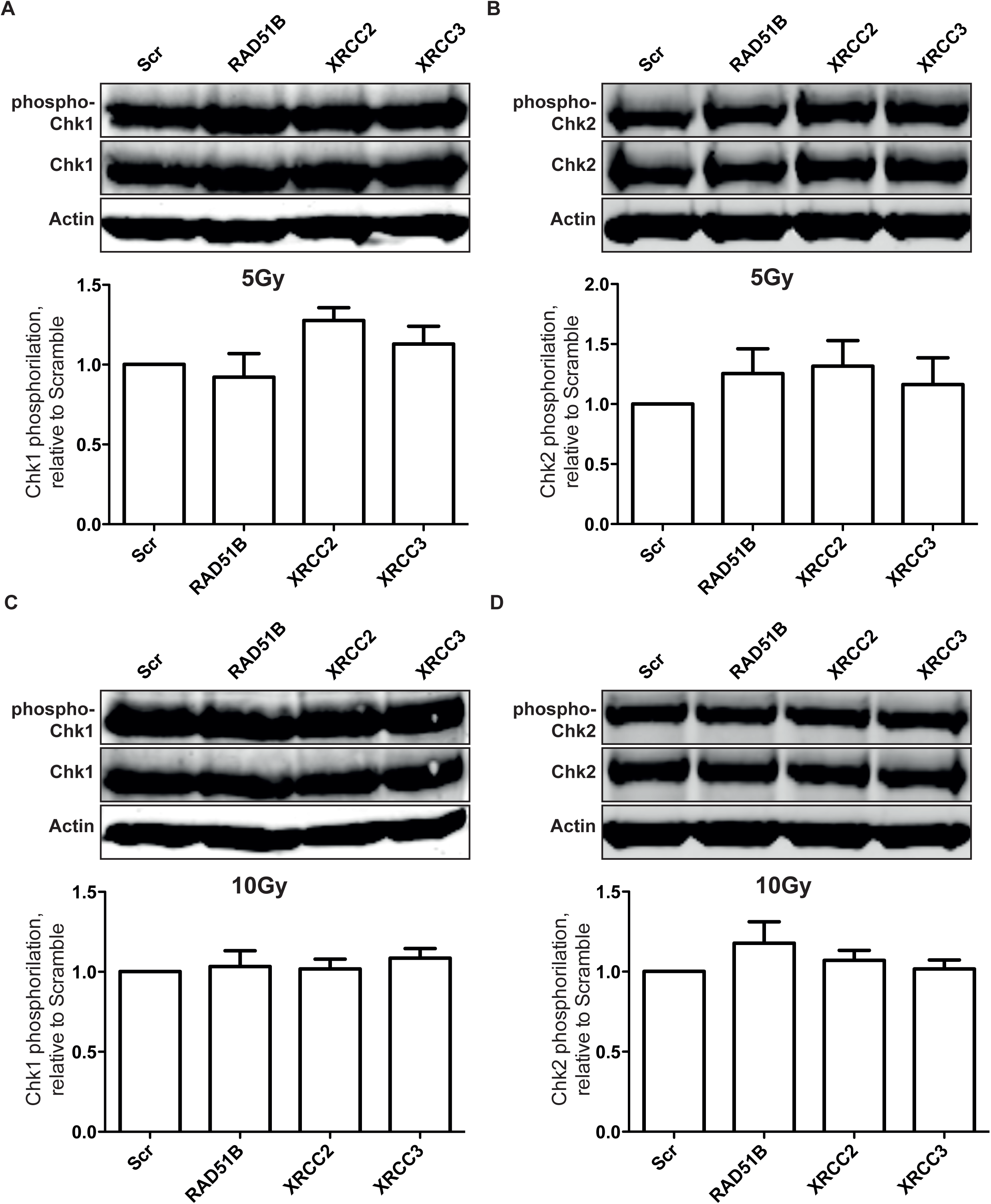
Checkpoint proficiency upon RAD51 paralogs depletion. A, ATR activation measured as CHK1 phosphorylation. Protein samples were obtained from cells infected with shRNAs against the indicated genes 1 h after treatment with 5Gy of ionizing radiation. After resolving the proteins in an SDS-PAGE, proteins were immunodetected with the indicated antibody. CHK1 phopshorylation proficiency was calculated as the ratio of the phosphorylated versus the total form of the protein and normalized with the scramble control. A representative western blot and the quantification of three independent experiments is shown. B, As in A, but for CHK2 threonine 68 phosphorylation. C, As in A but after 10 Gy of ionizing irradiation D, As in B but after 10 Gy of ionizing irradiation.

### Retention of NHEJ protein is affected by RAD51 paralogs

As we could not observe any major role of the RAD51 paralogs in early steps of HR, we wondered if they might be affecting NHEJ instead. In order to investigate such hypothesis, we measured the accumulation of Ku80 at the sites of DSBs upon reduction of RAD51 paralogs protein levels. Strikingly, we observed a mild increase in Ku80 retention at damaged DNA when RAD51B, XRCC2 or XRCC3 were depleted (Figure 4). Those data suggested that RAD51 paralogs interfered with NHEJ factor retention. Moreover, they indicate that it is possible to affect the NHEJ/HR ratio by affecting NHEJ, without altering DNA-end resection.

**Figure 4.**
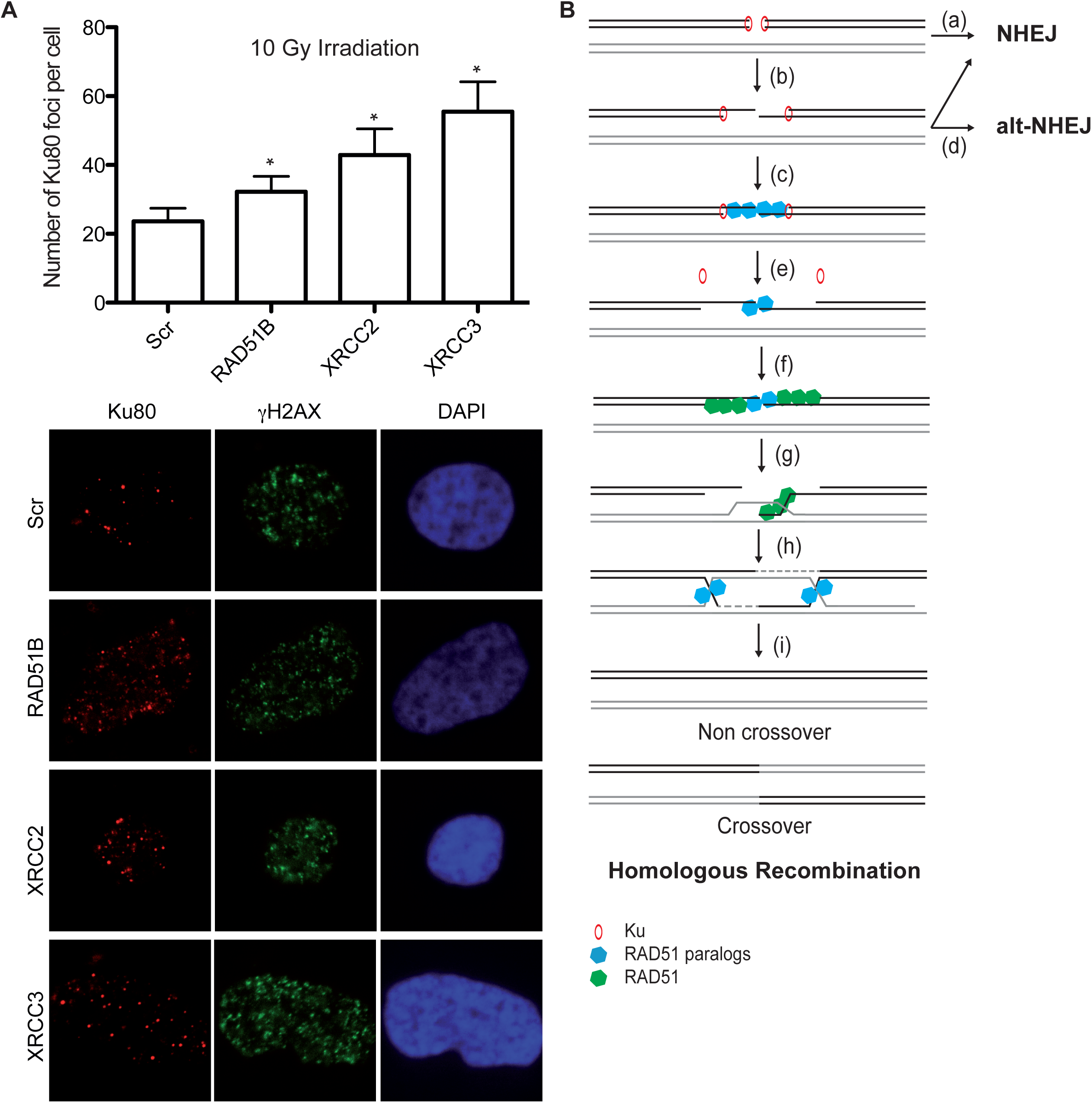
Ku80 retention in cells downregulated of RAD51B, XRCC2 and XRCC3. A, Cells depleted of the indicated genes were irradiated (10 Gy) and let to recover for 5 min. Then, samples were immunostained with anti-Ku80 antibody. A representative image upon depletion of each protein is shown. The number of Ku80 foci was calculated using metamorph as described in the methods section. At least 100 cells were scored and the median calculated. The graph represents the average of the median and the standard deviation of three independent experiments. B, Schematic representation of RAD51 paralogs roles during recombination. Upon the appearance of a break on the DNA Ku is immediately bound to the DNA ends. NHEJ can then take place (a) or limited resection could occur and Ku can be kept on the DNA (b). In the absence of RAD51 paralogs the presence of Ku can still promote NHEJ (d). Only when the paralogs are bound to ssDNA Ku is completely displaces (e), and HR takes place. RAD51 paralogs facilitate other HR steps such as strand invasion (f-g) or HJ resolution (h-i).

## Discussion

RAD51 paralogs are well-established factors involved in the repair of broken DNA at the level of RAD51-filament formation/stability and HJ metabolism (12, 14, 17, 24, 29, 43). Here, we show that they have roles at even earlier events, i.e. the decision between HR and NHEJ. This novel function can contribute to the decrease in overall recombination observed upon the paralogs depletion. Moreover, it can explain why the paralogs have been related with the RAD51-independent recombination SSA in *Arabidopsis thaliana* (44). This new function is shared by all five RAD51 paralogs, as seen for other recombination roles (44, 45), and it is independent of RAD51 itself, as RAD51 downregulation do not share such phenotype.

The decision to repair a broken DNA molecule by NHEJ or HR is tightly regulated. So far, the best know regulatory step was DNA-end resection (6, 26). The formation of RPA-coated ssDNA that follows DNA-end processing was considered enough to block the retention of the Ku70-Ku80 heterodimer, hence forcing the break to be repaired by HR (6, 26). Even the lack of Ku proteins was related to an increase on HR due to the unprotecting of the break to the resection machinery (46). However, our data suggest a resection independent role of Rad51 paralogues in controlling the balance between HR and NHEJ. We propose that the main role of RAD51 paralogs in repair pathway choice might occur after DNA end resection initiation, and might reflect resection extension (Figure 4B). RAD51 paralogs, as RAD51 itself, have the ability to bind double stranded and single stranded DNA (40). Indeed, in agreement with them modulating the ratio between NHEJ and HR after DNA-end resection, they bind ssDNA better than dsDNA (40) and they are recruited to damaged DNA post-resection in an RPA-dependent manner (25). In fact, recruitment to ssDNA stimulates the ATPase activity of RAD51 paralogs (40). One possible scenario is that short-range resection, at a level that does not interfere with the intrinsic DNA binding activity of Ku dimer (47), might be sufficient to recruit the RAD51 paralogs (Figure 4B, b-c). Indeed, Ku binds to artificial molecules with single-to-double stranded DNA transitions such as gaps or bubbles (48, 49). This recruitment of the paralogs to limited ssDNA would promote the local activation of their ATPase activity, somehow facilitating the displacement of the Ku heterodimer from the ssDNA-dsDNA transitions (Figure 4B, e). This crosstalk between ssDNA, RAD51 paralogs and the Ku dimer could provide a fail-safe mechanism for DSB repair. Thus, after initiation of short-range resection by CtIP and the MRN complex, Ku will stay bound to the DNA (Figure 4B, b). If those limited-processed breaks could not recruit RAD51 paralogs, i.e. cannot be efficiently engaged in HR, the Ku complex still present on the DNA could catalyze NHEJ. As DNA would have been partially resected, such repair will not be, probably, error-free, but might require an alternative, mutagenic NHEJ pathway (Figure 4B, d). This RAD51-independent role might explain why the absence of several paralogs change the pattern of repair during Ig diversification from HR-dependent gene-conversion to hyper-mutation (50). Only when RAD51 paralogs are recruited to DNA, anticipating the formation of the RAD51 presynaptic complex, the Ku complex is removed from the vicinity of the break, effectively blocking NHEJ and forcing the damage to be repaired by HR (Figure 4B, e). Although we could not exclude a direct interaction between Ku heterodimer and RAD51 paralogs, additional and still not characterized proteins might mediate such effect. Along those lines, it has been shown in *Saccharomyces cerevisiae* that Rad55 and Rad57, the yeast RAD51 paralogs, counteract the antirecombinogenic helicase Srs2 (52) and that Srs2 and Yku70 collaborate during NHEJ (53). Moreover, common interactors of both Ku and the RAD51 paralogs are known (54–56). Only after Ku removal, longer resection might take place, explaining the small but consistent reduction on the length of resected DNA we observed using SMART, mainly upon XRCC2 depletion. In fact, we cannot completely exclude a subtle role of the Rad51 paralogues in DNA end resection. Indeed, a connection between resection and the Rad51 paralogues has been found in rice, in which COM1 (CtIP plant homologue) recruitment to meiotic DSB requires XRCC3 (51). However, the resection defect observed upon XRCC2 downregulation is too small to account for the unbalance observed with the SSR. Finally, RAD51 paralogs will facilitate recombination by acting at the level of strand invasion together with RAD51 (Figure 4B, f-g) and HJ resolution (Figure 4B h-i).

## Methods

### Cell culture, lentiviral infection, and cell survival

U2OS cells were grown in Dulbecco’s modified Eagle’s medium (Sigma-Aldrich) supplemented with 10% fetal bovine serum (Sigma-Aldrich), 100 units/ml penicillin, 100 µg/ml streptomycin (Gibco) and 2 mM L-glutamine (Sigma-Aldrich). Lentiviral particles containing the different shRNAs were obtained as previously described (28). Cells containing plasmids were selected and maintained in culture adding 1 µg/ml Puromycine (Sigma-Aldrich). Information about the shRNA used in this paper can be found in the supplementary information (Table S1).

### Flow citometry (FACS)

Cells were harvested, washed with PBS and resuspended with ice-cold PBS. 70% EtOH was added dropwise while vortexing at low speed and then fixed at 4°C 2 hours. Cells were washed with PBS and treated with 20 mg/ml RNAse A (Sigma-Aldrich) and 1 mg/ml propidium iodide diluted in PBS (Fluka). Cells were incubated at 37°C for 30 min and analyzed. Data represent two experiments.

### Immunofluorescence microscopy

U2OS cells were infected with lentivirus harboring the indicated shRNA targeted against RAD51 paralogs or a control sequence. After 96 h, cells were irradiated (10 Gy), incubated for foci formation and then collected. Different protocols were used as described before for RPA (41), RIF1 (39) and Ku80 (57)foci detection. RPA foci formation was scored as the percentage of cells that have RPA foci from the total number of cells. The median number of RIF1 and Ku80 foci per cell was calculated using the software Metamorph as the number of dots present in the nucleus (defined by DAPI) on those cells that show H2AX staining. All data represent the average of a minimum of three experiments.

### NHEJ/HR balance analysis

U2OS cells bearing a single copy integration of the reporter SSR (NHEJ/HR balance) were used to analyze the different DSB repair pathways as previously published (28). One day after seeding, they were infected with a lentivirus harboring an I-SceI and labeled with BFP (58) using a M.O.I (multiplicity of infection) of 5. A day after infection the same volume of fresh medium was changed. Cells were grown during 48 hours after I-SceI infection, fixed with 4% paraformaldehyde, stained with Hoechst and washed with PBS prior visualization with a FACs for blue, green and red fluorescence. The NHEJ/HR ratio was calculated taking into account green versus red cells in each condition as published (28). This ratio was relativized with the control shRNA (Scramble). Those conditions that skew the balance towards an increase in NHEJ repair result in fold increase below 1. Data represent the average of three independent experiments.

### Immunoblotting

Extracts were prepared in Laemmli buffer (4% SDS, 20% glycerol, 120 mM Tris-HCl, pH 6.8), and proteins were resolved by SDS-PAGE and transferred to PVDF (Millipore) followed by immunoblotting. Western blot analysis was carried out using the antibodies listed in Supplementary Tables S2 and S3. Results were visualized using an Odyssey Infrared Imaging System (Li-Cor). To quantify Chk1 and Chk2 phosphorylation, protein abundance was measured using the Li-Cor software, and the ratio between the phosphorylated species versus total protein was calculated. Those ratios were then relativized with respect to control cells expressing shScramble. Data represent the mean of four experiments.

### Single molecule analysis of resection tracks (SMART)

SMART was performed as previously described (42). Briefly, U2OS cells depleted of the indicated proteins were grown in the presence of 10 µM bromodeoxyuridine (BrdU, GE Healthcare) for 24 h. Cultures were then irradiated (10 Gy) and harvested 1 hour after. Cells were embedded in low-melting agarose (Bio-Rad) followed by DNA extraction. To stretch the DNA fibers, silanized coverslips (Genomic Vision) were dipped into the DNA solution for 15 min and pulled out a constant speed (250 µm/sec). Coverslips were baked for 2 h at 60°C and incubated directly without denaturation with an anti-BrdU mouse monoclonal (Table S2). After washing with PBS, coverslips were incubated with the secondary antibody (Table S3). Finally, coverslips were mounted with ProLong^®^ Gold Antifade Reagent (Molecular Probes) and stored at –20°C. DNA fibers were observed with Nikon NI-E microscope and PLAN FLOUR40× / 0.75 PHL DLL objective. The images were recorded and processed with NIS ELEMENTS Nikon software. For each experiment, at least 200 DNA fibers were analyzed, and the length of DNA fibers were measured with Adobe Photoshop CS4 Extended version 11.0 (Adobe Systems Incorporated). Data represent the average of the medians of three experiments.

### Statistical analysis

Statistical significance was determined with a paired t student test using the PRISM software (Graphpad Software Inc). Statistically significant difference was labeled with 1, 2 or three asterisk if p<0.05, p<0.01 or p<0.001 respectively.

## Acknowledgements

We want to thank members of the lab for helpful advice and Cristina Cepeda-García and Sonia Jimeno for critical reading of the manuscript. This work was funded by an ERC Starting Grant (DSBRECA). FM-N is funded by a FPU fellowship from the Spanish Ministry of Education.

## Author contributions

DG-C discovered that RAD51 paralogs depletion lead to an unbalance in the SSR system. FM-N performed all the others experiments described. The project was conceived, designed, and developed by PH with the help of FM-N. All authors contributed to the discussion of the results. PH wrote the paper, with the feedback of FM-N and DG-C.

