## Supplementary Materials for "RAD51 paralogs regulate double strand break repair pathway choice by limiting Ku complex retention"

### Supplementary legends

**Supplementary Figure S1: Reduction of the RNA levels upon shRNA-mediated depletion against Rad51 paralogs.** Quantitative RT-PCR data was measured for mRNA in cells stably transfected with the indicated shRNAs. The mRNA level of each gene was normalized to actin mRNA. The mRNA level of each indicated gene in a cell line infected with the shRNA targeted against such gene (white bars) was normalized to the levels in a cell line stably transfected with an shRNA with a scrambled sequence (Black bars). Average and s.e.m. of a minimum of three experiments with triplicates is plotted.

### Supplementary methods

#### Quantitative PCR

$10^6$ U2OS cells were harvested in parallel with each experiment to check depletion of the RAD51 paralogs. For RNA extraction, RNeasy Mini Kit (Qiagen) was used using the manufacturer instructions. Then cDNA was obtained by QuantiTect Reverse Transcription Kit (Qiagen) following the kit instructions. For quantitative PCR, samples were mixed with iTaq Universal SYBR Green Supermix (BioRad) and the corresponding pair of primers (Supplementary Table S4). 96 well plates were run in a 7500 Fast Real-Time PCR System (Applied Biosystems) in Fast mode. Each experiment data was obtained from experimental triplicates.

### Supplementary tables

**Table S1: shRNAs used in this study**

| Target gene | Description | Supplier |
|---|---|---|
| **Rad51** | TRCN0000018877 | Sigma |
| **RAD51B** | TRCN0000151962 | Sigma |
| **RAD51C** | TRCN0000330043 | Sigma |
| **RAD51D** | TRCN0000151019 | Sigma |
| **XRCC2** | TRCN0000255930 | Sigma |
| **XRCC3** | TRCN0000083043 | Sigma |
| **CtIP** | TRCN0000318736 | Sigma |
| **Scramble** | SHC016 | Sigma |

**Table S2: PrimaryAntibodies used in this study.** IF, immunofluorescence. WB Western blotting. SMART, Single Molecule Analysis of Resection Tracks

| Target protein | Application | Supplier | Reference |
|---|---|---|---|
| **RPA32** | IF | Abcam | ab2175 |
| **RIF1** | IF | Bethyl Laboratories | A300-568A |
| **Ku80** | IF | Thermo Scientific | MA1-23314 |
| **γ-H2AX** | IF | Cell Signaling | 2577L |
| **γ-H2AX** | IF | Millipore | 05-636 |
| **Chk1 phospho S345** | WB | Cell Signalling | 2348 |
| **Chk1** | WB | Santa Cruz | Sc-8408 |
| **Chk2 phospho T68** | WB | Millipore | ABE33 |
| **Chk2** | WB | Millipore | 05-649 |
| **β-Actina** | WB | Abcam | Ab8227 |
| **BrdU** | SMART | Sigma | B5002-100MG |

**Table S3: Secondary antibodies used in this study.** IF, immunofluorescence. WB Western blotting. SMART, Single Molecule Analysis of Resection Tracks

| Antibody | Source | Application |
|---|---|---|
| **Alexa Fluor 594 goat anti-mouse** | Invitrogen | IF, SMART |
| **Alexa Fluor 488 goat anti-rabbit** | Invitrogen | IF |
| **IR-Dye 680 RD goat anti-mouse IgG (H+L)** | Li-Cor | WB |
| **IR-Dye 800CW goat anti-rabbit IgG (H+L)** | Li-Cor | WB |

**Table S4: Oligonucleotides used in this study.**

| Primer name | Sequence |
|---|---|
| **F-ACTB qPCR** | ACGAGGCCCAGAGCAAGA |
| **R-ACTB qPCR** | GACGATGCCGTGCTCGAT |
| **F-CtIP qPCR** | AGAAATTGGCTTCCTGCTCAAG |
| **R-CtIP qPCR** | GAAAACCAACTTCCCAAAAATTCTC |
| **F-RAD51 qPCR** | GGTGAAGGAAAGGCCATGTA |
| **R-RAD51 qPCR** | TATCCAGGACATCACTGCCA |
| **F-RAD51B qPCR** | AAGAGAGGCATCCTCCTTGAA |
| **R-RAD51B qPCR** | GCTCCACTCAGATGGGTTGT |
| **F-RAD51C qPCR** | AAACCCTCCGAGCTTAGCA |
| **R-RAD51C qPCR** | TGTGACTCAGATGTACCAGCA |
| **F-RAD51D qPCR** | TCCACGAGGAACTGAAGACC |
| **R-Rad51D qPCR** | ACCTGGGCCTCCTACAATTT |
| **F-XRCC2 qPCR** | GGCGGGAAAGTTGAGTCTCT |
| **R-XRCC2 qPCR** | CTACCTTCAAGTCGGGCAAG |
| **F-XRCC3 qPCR** | CAGCTCGGCAGGGAAGAC |
| **R-XRCC3 qPCR** | CGTGCAGATGTAGACGGC |

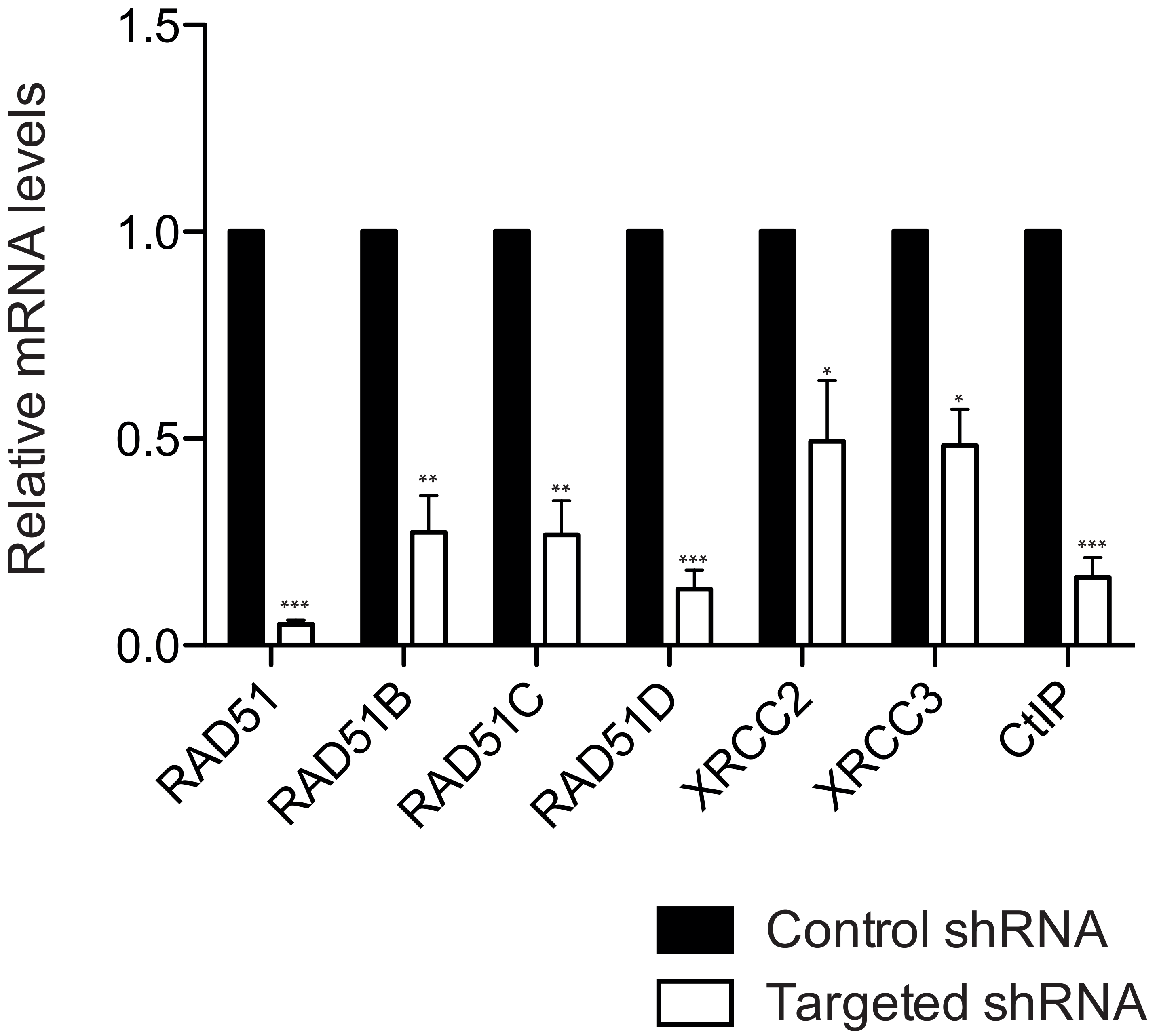


Mejías-Navarro et al Supplementary Figure 1
